# Bridging species boundaries: eDNA and genetic analysis reveal hybrid settlement at the very end of American eel distribution

**DOI:** 10.1101/2024.12.05.626534

**Authors:** Magnus Wulff Jacobsen, Rasmus Nygaard, Jens Frankowski, Aja Noersgaard Buur Tengstedt, Julius Nielsen, Michael M. Hansen, Rasmus Hedeholm, Anja Retzel, Ida Hedal, Paulina Urban, Casper Gundelund Jørgensen, Rasmus Stenbak Larsen, Sara Maggini, Peter Rask, Einar Eg Nielsen

## Abstract

Documenting species distributions and hybridization patterns is paramount for elucidating biogeography and understanding speciation processes. Here we combined genetic specimen analysis and environmental DNA (eDNA) to investigate the presence of American eel (*Anguilla rotrata*) and American × European eel (*Anguilla anguilla*) hybrids in Greenland freshwater. We further tested the use of eDNA to document hybridization by using European eel mtDNA as a proxy for hybrid occurrence. Overall, eDNA analysis detected mainly American eel but also European eel mtDNA. This finding was validated by DNA sequencing, which identified 3 out of 26 captured eels (14.3%) carrying European eel mtDNA. Five eels (19.2%), including all three with European mtDNA, were heterozygous for species-specific nuclear gene variants, supporting a hybrid ancestry. Further, eDNA successfully identified eels in lakes where they were caught by fyke net fishing, extending their confirmed northern range by 40 km, and indicated eel presence >200km further north. The study provides an empirical demonstration of the use of eDNA to document hybrid occurrence and extends the reported northern distribution of eels in Greenland significantly beyond previous observations. Further, the existence of hybrid eels in Greenland may be key for understanding the complex mechanisms of hybridization between American and European eels.

## Introduction

Documenting species occurrences and distributions is paramount for elucidating species biology and ecology and is crucial for effective management and conservation efforts^1^. In cases where species are hybridizing, information about hybrid occurrence and hybridization patterns may further hold important insights for understanding the mechanisms maintaining species boundaries^2,3^.

The catadromous European eel (*Anguilla anguilla*) and American eel (*Anguilla rostrata*) are categorized as respectively critically endangered and endangered by the IUCN^4,5,6^. Both species spawn in the frontal zones of the Sargasso Sea^7^, are panmictic^8,9,10,11^ and show a widespread distribution ranging from the subtropics to the Arctic^12^. The American eel has its reported northernmost occurrence in southwestern Greenland while European eel has its northernmost distribution in Iceland and Northern Norway^12^. European and American eels can hybridize^13,14,15,16^ and genomic studies indicate low levels of gene flow between the species^8^. Notably, the occurrence of hybrids is rare and observations of first-generation hybrids (F1-hybrids), as well as first- and second-generation back-crosses, have almost exclusively been documented in Iceland^13,14,16^, with a frequency of hybrids >10%^13,16^. While the reason behind this phenomenon remains unknown, it is potentially linked to an intermediate larval dispersal distance in F1-hybrids and immediate backcrosses, leading them to specifically settle in Iceland, which is approximately midway between the American and European continents^14^. Interestingly, eels in Iceland are predominantly either pure European eel or backcrosses in the direction of European eel, and F1- and later hybrid classes, almost exclusively carry European eel mtDNA^13,14,16^. This has led to the suggestion that gene flow between species is primarily unidirectional^17,18^. However, genome scale analyses of the two species support a scenario of bidirectional gene flow^19^. This points to the intriguing possibility that admixed individuals in the direction of American eel settle at other yet undiscovered localities.

The hitherto reported northernmost locality in in the world^12^ with American eel is the Greenlandic lake in Faeringehavn situated approximately 80 km south of the capital of Nuuk^20^. Previous species identifications relied on vertebrae counts, which are not 100% diagnostic, as they overlap with European eel^12^. Hence, it is currently unknown whether hybrids may exist in Greenland. Currently, American eel has only been documented from a handful of lakes in Greenland^20^, despite a potentially wider distribution across numerous lakes in Greenland. However, documenting fish distributions in remote Arctic regions is limited by the large logistical challenges and the absence of tradition for fishing with traps and nets in freshwater. Environmental DNA (eDNA) analysis represents an effective, non-invasive and cost-effective method for biodiversity monitoring^21,22,23^, including in Arctic regions^24,25,26^. This approach offers ease of sampling and high detection sensitivity, providing a suitable alternative or supplement to conventional direct fishing methods^21,22,23,24^. Previous research supports the possibility to use of eDNA for documenting occurrence of European^27,28^ and American eel^29,30^. However, despite the potential application no studies have used eDNA for monitoring eels in the Arctic. Further, the applications of eDNA are expanding with more research showing a potential use of the method to generate population genetic data^31,32^. For example, several papers have shown how eDNA can be used for population genetic inferences by analyzing mtDNA haplotype diversity in fish^33,34^, sharks^35,36^ and whales^37,38^. Environmental DNA has also been suggested as a tool for investigating hybrid occurrence^31^ in aquatic systems. However, while theoretically possible, empirical validation is lacking. This is likely due to the predominantly use of maternally inherited mtDNA, making eDNA well suited for assessing species occurrence and contact boundaries, but not for analyzing admixture, which require additional nuclear data^39^. One exception might be the European and American eels. They show distinct species boundaries^12^ and all documented cases of individuals carrying mtDNA from the other species have been designated as hybrids^14,15^. This missing introgression is potentially attributed to cytonuclear incompatibility, related to genes within the ATP synthase complex causing mismatches and protein dysfunction in later generations of hybrids^40,41,42^. Thus, in Atlantic eels, findings of mtDNA mixtures likely serve as a proxy for hybrid occurrence, and thus eDNA could be used for identification of areas of likely hybrid occurrence.

In this study, we analyzed for both American and European eel eDNA and examined captured specimens using genomic and genetic markers to screen for potential hybrids. This approach allowed us to assess the use of eDNA to document hybrid eel occurrences and to determine whether hybrid eels settle in Arctic Greenland. We also applied environmental DNA without parallel fyke net fishing to explore the broader distribution of eels. Finally, we compare the use of eDNA and fyke net fishing to investigate the potential future use of DNA-based monitoring to map the distribution of American eels in Greenland.

## Methods

### Environmental DNA sampling and fyke net fishing

Sampling of environmental DNA was conducted from 22^th^-30^th^ of July 2022 from (N=16 freshwater localities within the Nuuk Fjord (Nuup Kangerlua), as well as south of the fjord including the lake in Faeringehavn (Fig. 1). All sites included lakes with river access to the sea and most samples (N=14) were collected immediately downstream where the river left the lake, while two were collected further downstream (Table 1). Additional four localities were sampled from western Greenland (i.e., Kangerlussuaq and Sisimiut) and eastern Greenland (Kulusuk city lake and Nerlerit airport lake). The samples from Sisimiut were collected directly from the river outlet in the fjord (Fig 1).

**Table 1.** Fishing and qPCR results for each eDNA sampling location. Number of positive qPCR triplicates >LOD and number of overall positive qPCRs are noted for each sample replicate (a-c) with detected eel eDNA. Concentrations (copies/L) are calculated based on rounded copy estimates for the analyzed reactions. Since all estimates were below LOQ the shown values should be interpreted with caution. Fyke net fishing observations, catches and year of fishing are shown for the respective localities. NA = not available, nd. = no detections.

| Sample name | Location | Netfishing | No. Samples | Volume (L) | European eel qPCR |  |  |  | American eel qPCR |  |  |  |
| --- | --- | --- | --- | --- | --- | --- | --- | --- | --- | --- | --- | --- |
|  |  |  |  |  | a | b | c | ~Copies/L | a | b | c | ~Copies/L |
| Sisimiut | fjord | NA | 3 | 1 | 0/1 | 0/1 | 0/1 | 44.4 | nd. | nd. | nd. | 0 |
| Kangerlussuaq | lake outlet | NA | 3 | 2 | nd. | nd. | nd. | 0 | nd. | nd. | nd. | 0 |
| Taseraarsuk | river outlet | NA | 3 | 1 | nd. | nd. | nd. | 0 | nd. | nd. | nd. | 0 |
| Quagssugaq | river outlet | NA | 2 | 1 | nd. | nd. | nd. | 0 | nd. | nd. | nd. | 0 |
| Kapisillit 1 | lake outlet | no (2011) | 3 | 1 | nd. | nd. | nd. | 0 | nd. | nd. | nd. | 0 |
| Kapisillit 2 | lake outlet | no (2011) | 3 | 1 | nd. | nd. | nd. | 0 | nd. | nd. | nd. | 0 |
| Kapisillit 3 | lake outlet | NA | 3 | 1 | nd. | nd. | nd. | 0 | nd. | nd. | nd. | 0 |
| Nuuk city lake | lake outlet | NA | 3 | 0.6 | nd. | nd. | nd. | 0 | nd. | nd. | nd. | 0 |
| Kobbefjord | lake outlet | NA | 3 | 1 | nd. | nd. | nd. | 0 | nd. | nd. | nd. | 0 |
| Kangerdluk | lake outlet | NA | 3 | 1 | nd. | nd. | nd. | 0 | nd. | nd. | nd. | 0 |
| Qurajat2 | lake 2 outlet | 1 eel (2022) | 3 | 1 | nd. | nd. | 2/2 | 55.6 | nd. | nd. | nd. | 0 |
| Qurajat 1 | lake 1 outlet | no (2022) | 3 | 1, 1, 1.1 | nd. | nd. | nd. | 0 | nd. | nd. | nd. | 0 |
| Kigdlut ilua | lake outlet | NA | 3 | 1 | nd. | nd. | nd. | 0 | nd. | nd. | nd. | 0 |
| Agpânguit iluat 2 | lake 2 outlet | no (2022) | 3 | 1 | nd. | nd. | nd. | 0 | nd. | nd. | nd. | 0 |
| Agpânguit iluat 1 | lake 1 outlet | NA | 3 | 1 | nd. | nd. | nd. | 0 | nd. | nd. | nd. | 0 |
| Faeringehavn 1 | upper lake outlet | NA | 3 | 0.65, 0.6, 0.6 | nd. | nd. | nd. | 0 | nd. | nd. | nd. | 0 |
| Faeringehavn 2 | main lake outlet | 4 eels (2020-22),<br>21 eels (2011-13) | 3 | 1 | 1/2 | 0/1 | nd. | 44.4 | nd. | 1/1 | 2/2 | 155.6 |
| Faeringehavn 3 | river outlet | NA | 3 | 1 | nd. | nd. | nd. | 0 | 3/3 | 2/2 | 2/2 | 455.6 |
| Lake near faeringehavn | lake outlet | NA | 3 | 0.6 | nd. | nd. | nd. | 0 | nd. | nd. | nd. | 0 |
| Nerlerit airport lake | lake outlet | NA | 3 | 0.5 | nd. | nd. | nd. | 0 | nd. | nd. | nd. | 0 |
| Kulusuk city lake | lake outlet | NA | 3 | 0.5 | nd. | nd. | nd. | 0 | nd. | nd. | nd. | 0 |

**Fig 1.**
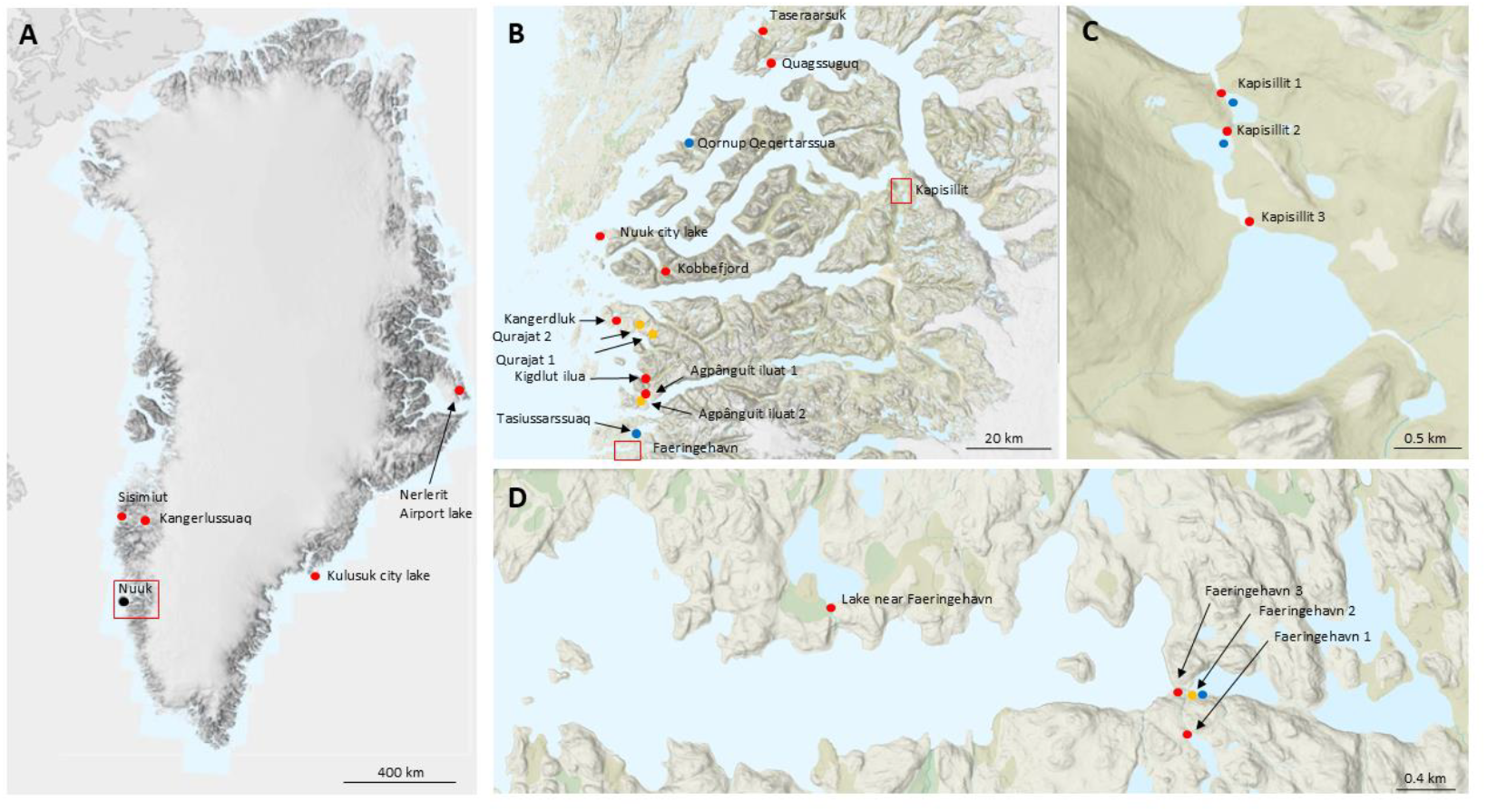
Sampling map showing all locations monitored by either eDNA (red), eDNA and net fishing (orange) and net fishing alone (blue). Localities in blue denote stations net fishing in 2011-2013.

From each locality we collected three ≤2L water replicates, which were filtered on site using a sterile 0.22 µm Sterivex filter (SVGPL10RC, Sigma-Aldrich, St. Louis, MO, USA). Quantitative PCR (qPCR) was performed to test for European eel^43^ and American eel^44^ eDNA and to estimate concentrations. All samples were analyzed in triplicate qPCR reactions with 8 µL template eDNA. Tests for potential PCR inhibitors were conducted by running an internal positive control (TaqMan™ Exogenous Internal Positive Control, Thermo Fisher Scientific). Reactions with inhibition observed as >1 ct-cycle increase for the IPC were rerun with 2 µL template eDNA. Species-specificity tests were conducted to exclude amplification of non-target species. LOD was estimated using the combined standard dilution series generated across plates (N=4) and calculated using the modelling approach described by Klymus et al.^45^. Additional metabarcoding analysis was performed on all samples from localities with positive qPCR detection of either American or European eel eDNA using the MiFish-U primer set targeting bony fishes^46^. Sequencing was performed on the Oxford nanopore MinION MK1C, using the R9.4.1 flow cell and associated library kit. The final sequences were demultiplexed and quality filtered (phred-score >10) with cutadapt^47^ and sequences clustered and transformed into consensus sequences using NGSpeciesID^48^ with a similarity score of 97 percent. Species identity was based on matching consensus sequences to a reference database using BLAST+ ver 2.12.0^49^ including all 5 known Greenland freshwater fish species and European eel eel (Supporting information, Note S1). Blast hits with >99% sequence similarity were used for species identity. For further information about qPCR and metabarcoding analyses, see supporting information Note S2.

Fyke net fishing was conducted from a subset of lakes in 2020-2022. This included the lake in Faeringehavn (fished in 2020-2022) and three more northernly located lakes with close access to the sea (Qurajat 1, Qurajat 2 and Anpângiut 2, all fished in 2022) (Fig.1 and Table 1). Fyke nets were baited with 5 kg of capelin (*Mallotus villosus)* and retrieved after approximately two weeks. Tissue samples from all eels (N=5) were collected by fin clips and stored in 96% ethanol.

### Extraction DNA sequencing of tissue samples

All tissue samples were extracted using the E.Z.N.A. Tissue DNA Extraction kit following the manufacturer’s instructions (OMEGA Bio-Tek, CA, USA). The mitochondrial cytochrome oxidase subunit I gene (*COI*) was sequenced using the primers (F1 and R2) described in Ward et al.^50^ and PCR products were Sanger sequenced using a SeqStudio Genetic Analyzer (Applied Biosystems). Sequences were trimmed using the Geneious Prime software (Geneious Prime 2022.0.1 https://www.geneious.com) and identified to species using NCBI Blast (https://blast.ncbi.nlm.nih.gov/Blast.cgi).

Full genome sequencing was performed for 4 individuals by DNBSEQ paired end (PE) sequencing (BGI, Hongkong, China) (one was not used due to DNA degradation because of nonoptimal storing conditions). To determine their genetic composition we included whole-genome sequence data from two European eels and two American eels used in a previous study^19^ and retrieved from the NCBI Sequence Read Archive (SRA accession no. <u>PRJNA554219</u>). The reads were aligned to the *Anguilla anguilla* genome assembly (GenBank accession no. <u>GCA_013347855.1</u>)^51^ using BWA MEM v.0.7.17^52,53^. Variants were called using the BCFtools pipeline^54^ and filtered using a range of software to ensure that only high quality SNPs were retained (see Supporting information Note S3 for details). Clustering of individuals was analyzed using ADMIXTURE v.1.3.0^55^ with 10-fold cross-validation and 100 bootstrap replicates for a value of *K* = 2.

DNA from an additional 21 eels collected in 2011-2013 from the main lake at Faeringehavn were analyzed. Cytogenetic genotyping was conducted after Frankowski & Bastrop^56^. Fragments of mitochondrial *CYTB* and the nuclear *18S* rRNA, freshwater opsin and deep sea opsin genes were Sanger sequenced to identify individuals with hybrids ancestry by comparing genotypes to American and European eels sampled on continental America or Europe (see Supporting information Note S4 for additional information).

## Results

### Fyke net fishing

A total of 25 eels were caught in the lake at Faeringehavn with 21 between 2011-2013, one in 2020 and three in 2021. One eel was caught in 2022 in a lake located ca. 40 km north of Faeringehavn in Qurajat (Qurajat 1, (Fig. 1). Fyke net fishing in seven additional lakes failed to record the presence of eels. These lakes included the 1^st^ and 2^nd^ lake in Kapisilit, a lake on the Island of Qornup Qeqertarssua and Tasiussarssuaq just north of Faeringehavn fished in 2011 and Qurajat 2, and Anpângiut 2 fished in 2022 (Fig. 1, Table 1).

### Environmental DNA sampling and analysis

The filtered water volume varied between 0.6-2L. All controls (i.e., field blanks, extraction blanks, NTCs) were negative. Twenty-four samples showed initial reaction inhibition and were diluted before further qPCR analysis. Eleven of these were from Faeringehavn and included all field replicates collected from the main lake and river outlet (Supporting Note S5). Quantitative PCR limit of detection (LOD) for three replicates was estimated to ∼2 copies per reaction for both assays. Overall, qPCR confirmed low amounts of eDNA from American eels at two sites downstream of the lake at Faeringehavn with an estimated 155.6 and 455.6 copies/L per site. All other samples were negative. Weak detections of European eel eDNA were also observed from the lake in Faeringehavn in the lake outlet, as well as from Qurajat 1 and Sisimiut (Table 1). Metabarcoding supported the finding of both American and European eel mtDNA at Faringehavn and European eel eDNA at Sisimiut. On the contrary, the method did not detect any eel eDNA from Qurajat 1. In addition, metabarcoding confirmed the presence of Arctic char (*Salvelinus alpinus*) and three-spined stickleback (*Gasterosteus aculeatus*) at all three analyzed localities (Fig 2). The overall proportion of European eel to American eel eDNA across all samples from Faeringehavn was estimated to 7.26%.

**Fig 2.**
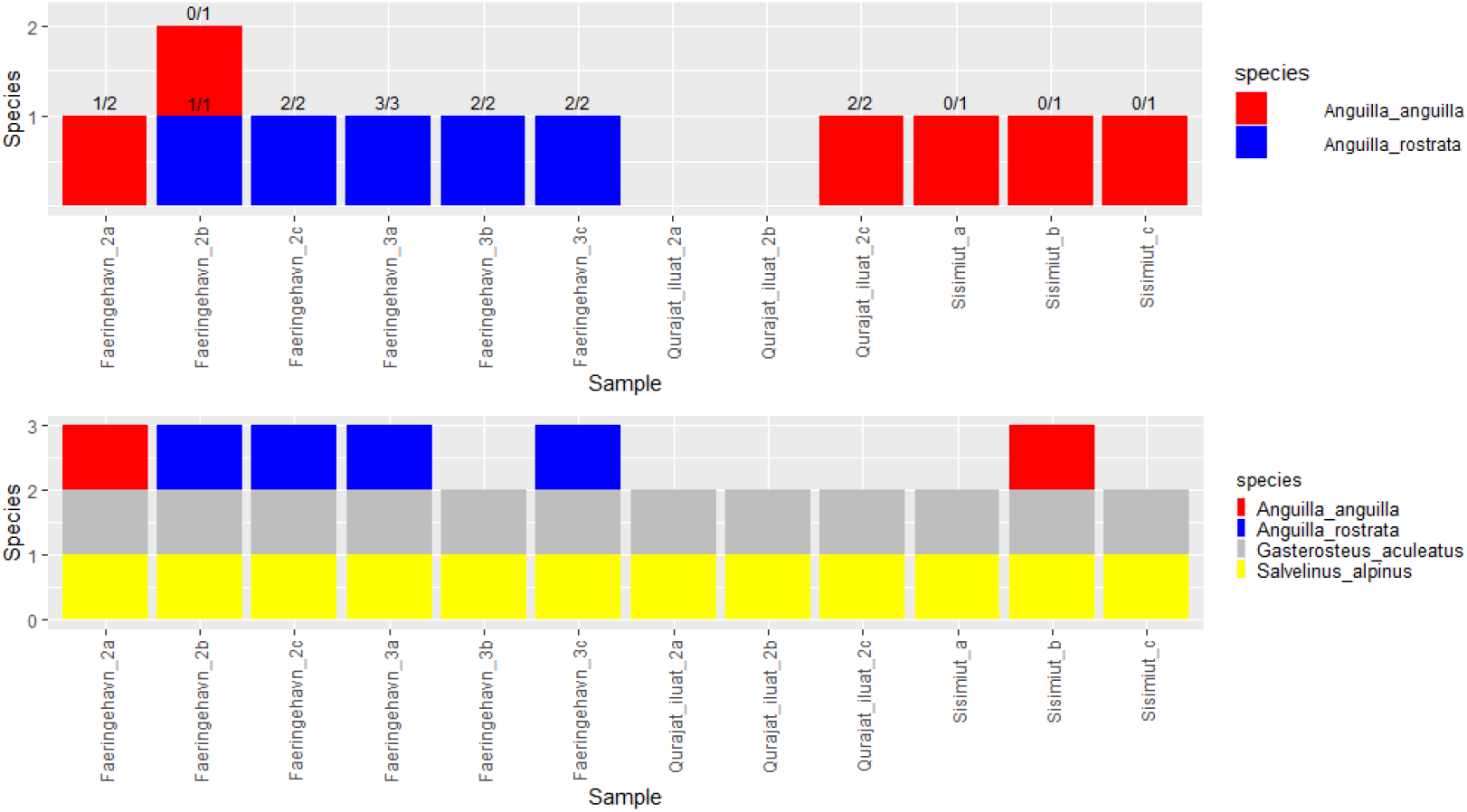
Overview of species detected using either qPCR (top) or metabarcoding (bottom). For the qPCR results, the numbers divided by the slash represent the number of triplicate analyses >LOD and total number with amplification. Only sampling stations with detected eel eDNA are shown.

### Genetic analysis of individuals

Mitochondrial sequencing of eels caught in 2022 confirmed that all five eels carried American mtDNA. Whole genome analysis of four of the individuals showed an American eel ancestry with no signs of hybridization (Supporting information Note S3), i.e., they were pure American eel. Of the 21 eels sampled in 2011 and 2012 in the lake in Faeringehavn, three had European mtDNA (14.3%), while the other carried the American mitogenome. To understand hybrid ancestry three nuclear genes were sequenced from these individuals and genotypes were compared to American and European eels sampled on continental America or Europe. All three eels with the European mtDNA and two with American mtDNA (23.8%) were heterozygote for the American and European 18S and opsin gene variants indicating that they were hybrids. The remaining eels were homozygote for the American variants (Table 2). The percentage of total analyzed specimens from Faeringehavn carrying European eel mtDNA was 12% and the percentage of hybrids was 20%.

**Table 2.**
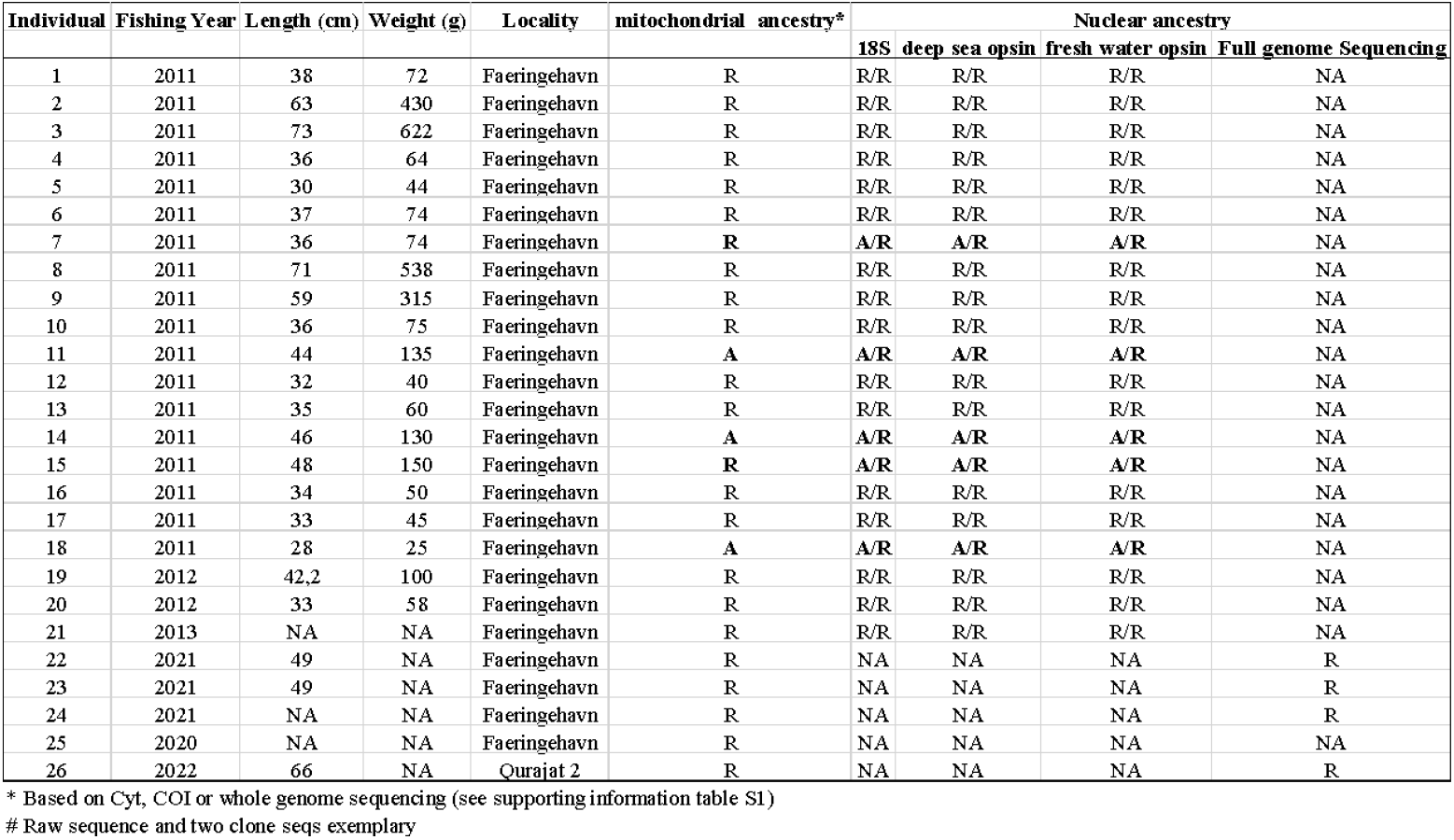
Information about fishing year, length (cm), weight (g), location, and genetic ancestry for all analyzed eels. R = *Anguilla rostrata*, A = *Anguilla anguilla*. For data accession numbers, see Supporting information Note S6. Individuals with nuclear heterozygote genotypes are shown in bold.

| Individual | Fishing Year | Length (cm) | Weight (g) | Locality | mitochondrial ancestry* | Nuclear ancestry |  |  |  |
| --- | --- | --- | --- | --- | --- | --- | --- | --- | --- |
|  |  |  |  |  |  | 18S | deep sea opsin | fresh water opsin | Full genome Sequencing |
| 1 | 2011 | 38 | 72 | Faeringehavn | R | R/R | R/R | R/R | NA |
| 2 | 2011 | 63 | 430 | Faeringehavn | R | R/R | R/R | R/R | NA |
| 3 | 2011 | 73 | 622 | Faeringehavn | R | R/R | R/R | R/R | NA |
| 4 | 2011 | 36 | 64 | Faeringehavn | R | R/R | R/R | R/R | NA |
| 5 | 2011 | 30 | 44 | Faeringehavn | R | R/R | R/R | R/R | NA |
| 6 | 2011 | 37 | 74 | Faeringehavn | R | R/R | R/R | R/R | NA |
| 7 | 2011 | 36 | 74 | Faeringehavn | R | A/R | A/R | A/R | NA |
| 8 | 2011 | 71 | 538 | Faeringehavn | R | R/R | R/R | R/R | NA |
| 9 | 2011 | 59 | 315 | Faeringehavn | R | R/R | R/R | R/R | NA |
| 10 | 2011 | 36 | 75 | Faeringehavn | R | R/R | R/R | R/R | NA |
| 11 | 2011 | 44 | 135 | Faeringehavn | A | A/R | A/R | A/R | NA |
| 12 | 2011 | 32 | 40 | Faeringehavn | R | R/R | R/R | R/R | NA |
| 13 | 2011 | 35 | 60 | Faeringehavn | R | R/R | R/R | R/R | NA |
| 14 | 2011 | 46 | 130 | Faeringehavn | A | A/R | A/R | A/R | NA |
| 15 | 2011 | 48 | 150 | Faeringehavn | R | A/R | A/R | A/R | NA |
| 16 | 2011 | 34 | 50 | Faeringehavn | R | R/R | R/R | R/R | NA |
| 17 | 2011 | 33 | 45 | Faeringehavn | R | R/R | R/R | R/R | NA |
| 18 | 2011 | 28 | 25 | Faeringehavn | A | A/R | A/R | A/R | NA |
| 19 | 2012 | 42.2 | 100 | Faeringehavn | R | R/R | R/R | R/R | NA |
| 20 | 2012 | 33 | 58 | Faeringehavn | R | R/R | R/R | R/R | NA |
| 21 | 2013 | NA | NA | Faeringehavn | R | R/R | R/R | R/R | NA |
| 22 | 2021 | 49 | NA | Faeringehavn | R | NA | NA | NA | R |
| 23 | 2021 | 49 | NA | Faeringehavn | R | NA | NA | NA | R |
| 24 | 2021 | NA | NA | Faeringehavn | R | NA | NA | NA | R |
| 25 | 2020 | NA | NA | Faeringehavn | R | NA | NA | NA | NA |
| 26 | 2022 | 66 | NA | Qurajat 2 | R | NA | NA | NA | R |

\* Based on Cyt, COI or whole genome sequencing (see supporting information table S1)

## Discussion

This is the first study to explore the wider distribution of freshwater eels in Greenland and represents the first genetic investigation of eels and hybrids in the region. It is further a pioneering study demonstrating the potential use of eDNA to analyze eel hybrid occurrence and the first study to show a high frequency of hybrid eels besides their documented occurrence in Iceland. Below we discuss the findings and their importance for understanding eel biology, ecology and evolution.

### Hybrid distribution and consequences for understanding introgression patterns in Atlantic eels

Full genome analysis of four eels uncovered that they all were pure American eels, aligning with the historical observations based on vertebrae counts conducted in the late-1900s^20^. Intriguingly, both eDNA and mtDNA sequencing of captured eel specimens revealed the presence of European eel mtDNA in Greenland eels. This has not been demonstrated before, but mirrors the situation in Iceland, where both species’ mtDNA have been reported^10,13,14,16,17^. Notably, eels in Iceland are predominantly genetically European^8,10,13,14,16,17^ with most individuals harboring European eel mtDNA^8,10,14,15,15,17^ (up to 3.5% of eels have shown to be American^14^). This sharply contrasts the predominantly American ancestry observed in Greenland eels in this study where only 14.3% of the captured eels contain the European mitogenome. Moreover, it is noteworthy that five eels sampled in 2011-2012 were heterozygous for the American and European 18S and opsin gene variants supporting a hybrid ancestry. Interestingly these eels included both hybrids with European and American mtDNA (respectively three and two of the 26 captured individuals; 11.5 and 7.7%). While the presented data precludes any conclusive inference on whether these individuals represent F1-hybrids or back crosses, it is likely that they constitute early-generation hybrids. Although speculative, the high proportion of hybrids with European eel mtDNA might indicate the existence of other localities with higher proportions of American eel hybrids. Nevertheless, with an observed proportion of 19.2% of specimens displaying genotypes consistent with early-generation hybrids, the observations from Greenland align with the range otherwise only reported in Iceland, which so far has shown the highest documented hybrid proportion of any known location (10.7% to 15.5%^16,17^). This implies an important role for Greenland in the settlement of hybrid eels and, consequently, for maintaining bidirectional gene flow between thetwo Atlantic eel species. Thus, future genomic analyses of these specimens could prove pivotal for elucidating how this unique system of hybridization is maintained, where hybridization occurs thousands of kilometers away in the Sargasso Sea^8,12,17^, Continental Europe and North America harbors almost exclusively pure European and American eel^8,9,10,11^, respectively, whereas Iceland and Greenland are nursery areas for hybrid eels. Hence, we suggest that both Iceland and Greenland act as sources of hybrids mediating further admixture in subsequent generations, with Icelandic eels introgressing in the direction of European eel, and Greenland eels introgressing into the American gene pool, leading to an overall pattern of bidirectional gene flow.

### Analyzing hybrid occurrence using environmental DNA

In this study, the genetic analysis of captured eels confirmed that all specimens with European eel mtDNA were hybrids, supporting the use of mtDNA as a proxy for hybrid presence. Our eDNA observations from Faeringehavn demonstrated that 7.26% of the detected eel mtDNA sequences belonged to European eel. The result matched well with the proportions observed for the captured specimens (12%), despite that these were mainly caught at another time point (2012-2013), suggesting stability of the observed genetic patterns. Thus, our study supports the use of eDNA as a good indicator for detection of hybrid eels and provides to our knowledge the first empirical demonstration of the use of eDNA for identification of potential hybrid occurrence in any species. Still, the use of mtDNA for hybrid detection in eel is challenged by the dependence on the presence of hybrids with European maternal ancestry. Hybrid documentation in cases where hybrids carry American mtDNA, which was also observed in this study, is only possible using nuclear DNA^39^. The application of nuclear eDNA should be investigated in the future, however, is challenged by the lower expected copy number of nuclear DNA compared with mtDNA^57^, leading to a potentially lower sensitivity and related need for collecting and analyzing much larger volumes of water than conducted here. However, the water in Greenland is very nutrient poor^58^ and has low turbidity rendering it feasible to filter larger volumes of water without problems. As such, Arctic freshwater systems like Greenland are likely ideal for testing this intriguing possibility.

### Distribution of freshwater eels in Greenland - insights from eDNA and fyke net fishing

Both qPCR results and fyke net fishing independently confirmed the presence of eels at Faeringehavn and Qurajat 1. The discovery of eels in Qurajat 1 extends the confirmed distribution of freshwater eels in Greenland ∼40 km north^12,20^. Notably qPCR exclusively detected European eel eDNA in Qurajat 1, while the captured individual carried American eel mtDNA. This outcome is likely attributed to chance, arising from low specimen density and mixed mtDNA ancestry of eels inhabiting the lake. This possibility finds support in the eDNA results from the main lake in Faeringehavn where European and American eel eDNA detections vary among samples, with some samples showing only presence of European eel DNA despite an overall higher concentration of American eel eDNA within in the lake (Table 1). European eel eDNA was also detected in Sisimiut suggesting that the distribution of eels in Greenland extends at least 200 km further north. This finding was further validated by the metabarcoding analysis, which confirmed the presence of European eel eDNA at Sisimiut and Faeringehavn, and therefore firmly excluded that these findings were not a result of unspecific amplification, for example of American eel. Whether the findings in Qurajat 1 and Sisimiut reflect European eel or hybrid eel eDNA cannot be verified from the study. However, the genetic findings from Faeringehavn showed that all specimens with European eel mtDNA were hybrids, which suggests that the findings of European eel eDNA in other Greenland lakes likely can be attributed to hybrid eel presence. Overall, this study supports the utility of employing eDNA as a cost- and time-effective alternative to fyke net fishing for mapping the distribution of eels in Arctic freshwater and supports previous conclusions regarding eDNA as an effective method for monitoring freshwater eels^27-30^. The information from eDNA, in turn, can guide traditional direct fishing efforts for complete genetic species confirmation. In regions with American and European hybrids, eDNA monitoring should involve analyzing for both American and European eel mtDNA using either qPCR or metabarcoding methods. The latter approach would additionally to documenting the presence of eels and eel hybrids, enable the collection of data on other species of fish such as Arctic char and three-spined stickleback as shown in this study.

### Eel occurrence in mid-western Greenland freshwater

Overall, both eDNA and fyke net fishing supported a low occurrence of eels in mid-Western Greenland, reaffirmingconclusion in and Jensen REF from 1937^59^ and in Böetius et al.^20^ from 1972 that eels are supposedly rare in Greenland. It is noteworthy that both fyke net fishing and eDNA supported a much higher density of eels in Faeringehavn than the other examined lakes. The river connecting the lake to the sea is short (only around 100 meters), with no large waterfalls or other barriers, which could impede migration of glass eels entering from the fjord. It is, however, also a characteristic shared with many of the other examined localities and therefore unlikely to be the main cause of the finding. As an example, Kapisillit lake system situated north of the capital Nuuk is a large lake system compromising >10 lakes in sequence separated by rivers^58^. Like Faeringehavn the system has easy access from the fjord. Due to the size of the system, it is significantly warmer than most other lake systems in the region^60,61^, which also likely explains the occurrence of Greenland’s only currently known self-sustaining Atlantic salmon population^58^. Hence, in many ways Kapisillit should be a perfect eel habitat, but neither fyke net nor eDNA supports any presence of eels in the system. Nielsen et al.^62^ also used eDNA and likewise failed to detect any eel eDNA in Kapisillit further supporting the results in this study. Kapisillit river is, however, located deep inside the Nuuk fjord and a migrating eel larvae will have met countless potential freshwater systems before reaching kapisillit river. Thus, a potential other explanation for the observations of high eel density in Faeringehavn may be that the lake is located closer to the open sea. As the eel larvae approach the continental shelf, they start metamorphosing into glass eels, and initiates migration into brackish or freshwater^12^. Here freshwater may act as a cue facilitating migration towards rivers and lakes^63^, which may explain that eels mainly end up in Faeringehavn. Interestingly, eDNA sample extractions from this locality were among the only ones to show initial reaction inhibition. This result could indicate the presence of higher concentrations of humic substances^64^, potentially linked to higher nutrient levels^65^, which could be another factor explaining higher eel numbers. Additional studies, incorporating measurements of physical and chemical parameters might help to test the above hypotheses.

## Ethics

Fishing was conducted following the acceptance from the Greenland government.

## Data accessibility

mtDNA and nuclear sequence data is available in Genbank (ref:xxxxxx). The genome data and metabarcoding data is available on xxx.

## Authors contribution

Author contributions: MWJ, JF, RN, RH and JN conceived and designed the study. RN, MWJ, JF, RSL, PU and CGJ conducted the fieldwork in 2020-2022, while JF, RN, RH, AR and PR conducted the fieldwork in 2011-2013. MWJ, IH and SM led the molecular and analytical eDNA work. MWJ analyzed the mtDNA data for eels collected in 2020-2022. JF analyzed the mtDNA and nuclear data for eels collected in 2011-2013. ANBT and MMH analyzed the full genome data. MWJ wrote the manuscript with contributions from ANBT, JF, EEN, MMH, RN, JN, PR, PU, SM, IH, RSL and CGJ.

## Conflict of interest declaration

We declare we have no conflict of interest.

## Funding

The work was funded by Elisabeth and Knud Petersen’s fund awarded to MWJ, RN and JN. Fieldwork in 2011-13 was supported by KIIN, European Fisheries Fond and the State of Mecklenburg-Western Pomerania

## Acknowledgement

We thank Britta Pedersen and Dorte Meldrup for laboratory assistance and Mala Broberg, Cornelia Albrecht, Manfred Stein and Kaj Sünksen for help during field work.

## Notes

### Competing Interest Statement

The authors have declared no competing interest.

