## Supporting information for "Bridging species boundaries: eDNA and genetic analysis reveal hybrid settlement at the very end of American eel distribution"

Rasmus Nygaard^2^

Jens Frankowski^3,4^

Aja Noersgaard Buur Tengstedt^5^

Julius Nielsen^2^

Michael M. Hansen^5^

Rasmus Hedeholm^2^

Anja Retzel^2^

Ida Hedal^1^

Paulina Urban^1^

Casper Gundelund Jørgensen^9^

Rasmus Stenbak Larsen^6^

Sara Maggini^7^

Peter Rask^8^

Einar Eg Nielsen^1^

^1^National Institute of Aquatic Resources, Technical University of Denmark, Vejlsøvej 39, DK-8600 Silkeborg, Denmark

^2^Greenland Institute of Natural Resources, Kivioq 2, P.O. Box 570, 3900 Nuuk, Greenland

^3^University of Rostock, Dept. Animal Physiology, Dr. Ralf Bastrop’s Lab, Rostock, Germany.

^4^Institute of Fisheries, State Research Centre for Agriculture and Fisheries Mecklenburg Western Pomerania, Rostock, Germany

^5^Department of Biology, Aarhus University, Ny Munkegade 114, DK-8000 Aarhus C, Denmark

^6^Department of Biology, University of Copenhagen, Ole Maaløes vej 5, DK-2200, Copenhagen N, Denmark

^7^Molecular Ecology and Evolution Bangor, School of Biological Sciences, Environment Centre Wales, Bangor University, Bangor, Gwynedd, LL57 2UW, UK

^8^Natural History Museum of Denmark, University of Copenhagen, Copenhagen, Denmark

^9^National Institute of Aquatic Resources, Technical University of Denmark, Kemitorvet 202, 2800 Kgs. Lyngby, Denmark

**Note S1**

**Database construction**

A database was established to include all five known species of Greenland freshwater fish, as well as European eel (*Anguilla anguilla*). The 5 species are Arctic char (*Salvelinus alpinus*), Atlantic salmon (*Salmo salar*), American eel (*Anguilla rostrata*), Three-spine stickleback (G*asterosteus aculeatus*) and the newly colonizing Pink salmon (*Oncorhynchus gorbuscha*). NBCI accession numbers on all sequences used in the database can be found in below table.

**Table S1. NCBI accession numbers for all sequences used in the database.**

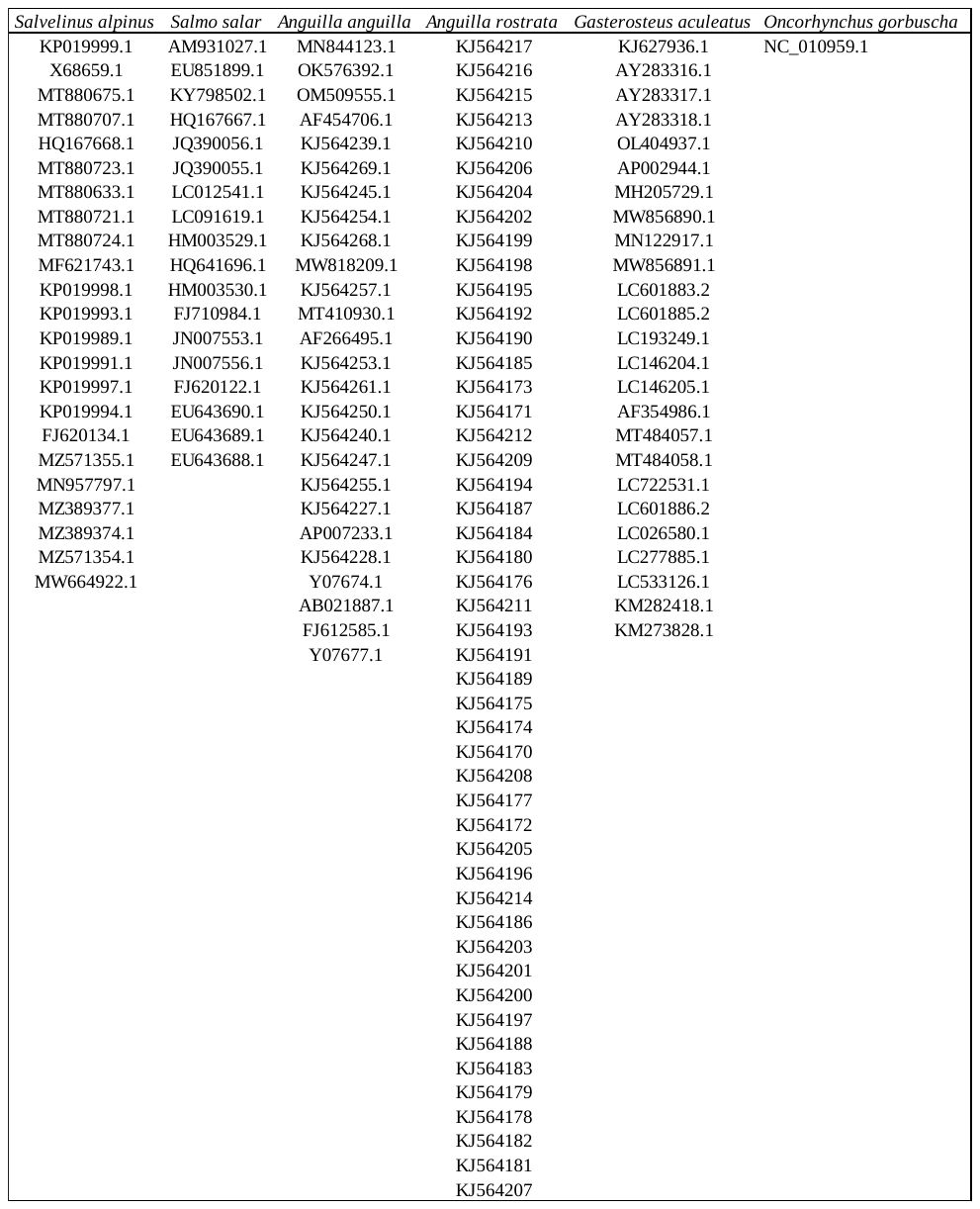

**Note S2**

**Laboratory processing of collected eDNA filters and results of eDNA analyses**

**Methods**

From each locality we collected three ≤1L water replicates, which were filtered on site using a sterile 0.22 µm Sterivex filter (SVGPL10RC, Sigma-Aldrich, St. Louis, MO, USA) and a vampire peristaltic sampler (Buerkle, Bad Bellingen, Germany). DNA preservation was obtained by passing 2 mL ATL buffer (Qiagen, Hilden, Germany) into the filters, which were subsequently closed and stored in a Styrofoam box until DNA extraction. DNA extraction was performed using the DNeasy blood and tissue extraction protocol and reagents (Qiagen, Hilden, Germany) and plant midi kit purification columns (Macherey-Nagel, Düren, Germany). DNA extractions included extraction blanks (i.e., extractions without DNA) to check for exogeneous contamination. Sample controls included NTCs (i.e. PCRs without template) and field blanks constituting either filtering of DNA-free nuclease free water or water collected at a low depth lake in the center of Nuuk where eels should not be present. After extraction, DNA concentration was measured using the Qubit flourometer and the Qubit dsDNA high sensitivity kit (ThermoFisher Scientific, Waltham, MA, USA).

Quantitative PCR (qPCR) was performed to test for American eel and European eel eDNA, as well as to estimate amplicon concentration. Assays developed for American eel^1^ and European eel were used^2^. Before eDNA analysis we performed a species species-specificity test by testing the assays on DNA from all known Greenland freshwater species (see Table S1). These samples included 1 ng DNA from three-spine stickleback, American eel, European eel, Atlantic salmon, Arctic char and pink salmon.

Quantitative PCR reactions were performed in 20 µL reaction volumes 8 µL of TaqMan Environmental Mastermix 2.0 (ThermoFisher Scientific, Waltham, MA, USA) and 2 µL template DNA. For the assay targeting American eel we used 500 nM forward primer, 500 nM reverse primer and 200 nM probe while the European eel assay used 800 nM forward primer, 1200 nM reverse primer and 300 nM probe. The probes were for both species Zen-probes. Reactions were run on a StepOnePlus Real-time PCR instrument (Life Technologies, Carlsbad, CA, USA), using 10 minutes initial denaturation at 95°C, followed by 50 cycles of 95°C for 15 seconds and 60°C for 30 seconds. Tests for potential PCR inhibitors were conducted by running an internal positive control (TaqMan™ Exogenous Internal Positive Control, Thermo Fisher Scientific) on all extracts using the same volume of extract as in the species tests. All samples were analyzed in triplicate qPCR reactions and compared to a standard dilution series using 1-1⨯10^5^ copies of gblock amplicons matching the target species. LOD was estimated using the combined runs of standard dilutions generated across plates (N=4) and calculated using the modelling approach described by Klymus et al.^3^.

Metabarcoding analysis was performed on all samples collected at the four localities with positive qPCR detection of either American or European eel eDNA. The localities included the main river and main lake at Faeringehavn, the lake outlet at Qurajat 1 and the river outlet sampled in Sisimiut. All samples were processed using the MiFish-U primer set targeting bony fish^4^. A total of three PCR reactions were conducted per sample. To discriminate between samples, PCR reactions were performed using different combinations of uniquely barcoded primers, so that each triplicate PCR held a unique genetic barcode. Each barcode was 7 bp long and had a minimum of 3 mismatches to other barcodes. Due to the relative high error rate introduced during MinION sequencing, barcodes were applied on both forward and reverse primers to ensure low risk of misidentification. The final PCR reactions were performed in 20 µL reaction volumes containing 10 µL of Amplitaq Gold Master Mix (Applied Biosystems, Foster City, CA, USA ), 0,16 µL BSA (20 μg/ µL), 5 μM of each primer and 2 µL of eDNA. Reactions were run on a standard PCR machine, using 10 minutes at 95°C, followed by 35 cycles including a 1 min DNA denaturation step at 94°C, a 1 min annealing step at 60°C and a 1 min extension step at 72°C. A final 5 min extension step at 72°C was performed after the 35 cycles. Following PCR, 5 µL from each triplicate PCR reaction was following pooled together and positive amplification was confirmed by gel electrophoresis. A volume of 5 µL from each of the pooled triplicates were finally pooled to make up the final sample for library construction. A DNA MinION library was prepared following the Nanopore protocol for Genomic DNA by Ligation using the SQK-LSK109 ligation kit (downloaded from Nanopore‘s webpage). Sequencing was conducted using a R9.4.1 flow cell and run on the Mk1C sequencing instrument enabling high quality base calling. The generated sequences were quality filtered (phred-score >10) and samples demultiplexed with cutadapt^5^ and sequences clustered and transformed into consensus sequences using NGSpeciesID^6^ with a similarity score of 97 percent. Species identity was based on matching consensus sequences to a reference database using BLAST+ ver 2.12.0^7^ including all 5 known Greenland freshwater fish species and European eel (Supporting information, Note S1). Blast hits with >99% sequence similarity was used for species identity.

**Results**

*qPCR analyses*

The two assays worked well and showed efficiencies and goodness-of-fit (R^2^) (as calculated from the standard dilution series) in the ranges considered good for quantitative analysis (R^2^ >0.99 and efficiency 90%<x<110%) (Bustin et al. 2009). The American and European eel assays did not cross amplify with DNA from any of the other known Greenland freshwater fish species. However, for both assays we observed minor cross-amplification with its sister-species (i.e., American or European eel). The difference in ct-values when analyzing of 1 ng of extracted DNA was approximately 16 ct-values for the assay targeting American eel and 10 ct-values for the assay targeting European eel (Table S1). Hence, while cross-amplification between the two eel species theoretically can lead to false-positives, such a result seems unlikely and should only occur in cases of extremely high eDNA concentration of the none target species, which generally seems unlikely. LOD for three replicates was estimated to 2.11 for European eel and 1.96 for American eel. LOQ was not calculated due to a high variability in ct-values for both assays, excluding estimation of LOQ based on the default CV variation threshold of 0.35. Instead a discrete approach was used, which estimated LOQ to 10 for both assays. An absolute LOD was tentatively set to a ct-value of 41 following Agersnap et al.^8^, which also tentatively matched the lowest ct-values observed from the standard dilution series (European eel 41.25 and American eel 40,4).

In general, most of the samples, as well as all field blanks, all extraction blanks and all NTCs were negative. Only a few samples showed amplification for American or European eel eDNA (for more details see main text).

**Table S2. Result of the cross-amplification tests. All test included triplicate qPCR tests and 1 ng of extracted DNA from target, and none target species.**

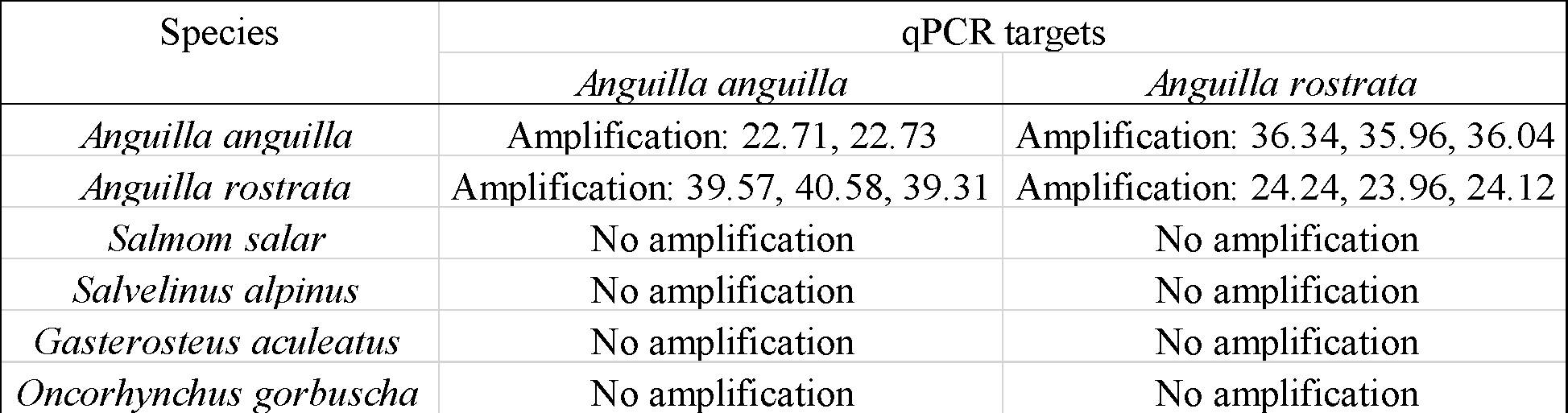

*Metabarcoding analysis*

After quality filtering and demultiplexing a total of 251,452 reads were retained. Following sequence clustering and database analysis a total of 169,223 sequences were left. The number of reads per sample differed amongst sampling stations with average numbers of 13,595.3 (SD: 4.181.9) for Qurajat 1, 12,650 (SD: 1.957.7) for Faeringehavn main lake outlet, 28,562.7 (SD: 6900.1) for Faeringehavn main river outlet, and 1,599.7 (SD: 704.8) for Sisimiut. The low number of sequences generated from the Sisimiut sampling stations matching the database were explained by a high number of sequences matching marine species (data not shown). This result was expected, as the samples were collected directly from the river outlet in the fjord. Metabarcoding supported the finding of both American and European eel mtDNA at Faringehavn and European eel eDNA at Sisimiut. On the contrary, the method did not detect any eel eDNA from Qurajat 1. In addition, metabarcoding confirmed the presence of Arctic char (*Salvelinus alpinus*) and three-spine stickleback (*Gasterosteus aculeatus*) at all three analyzed localities.

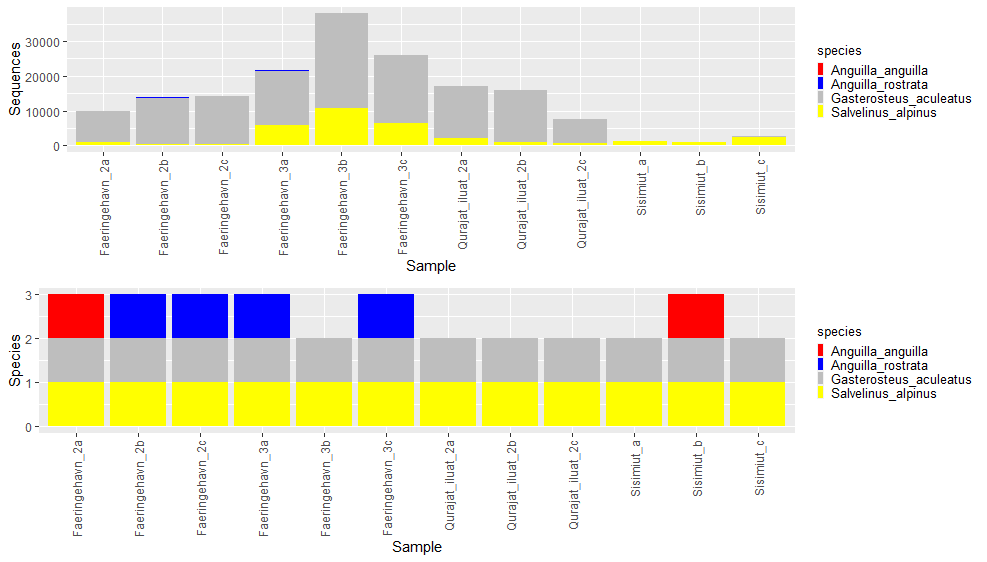

Fig S1. Metabarcoding results from each of the analyzed samples. Each sampling station includes three field replicates. The top figure shows the total number of sequences analyzed per sample and their species match. The bottom figure shows the species detected for each of the analyzed samples.

**Note S3**

**Whole genome analyses of four eels**

**Methods**

To determine the genetic composition of the four individuals sampled in Greenland, we included whole-genome sequence data from two European eels and two American eels used in a previous study (Nikolic et al., 2020) and retrieved from the NCBI Sequence Read Archive (SRA accession no. [PRJNA554219](https://www.ncbi.nlm.nih.gov/sra/?term=PRJNA554219)). Using BWA MEM v.0.7.17 (Li, 2013; Li & Durbin, 2009a) with default parameters, the reads were mapped to the *Anguilla anguilla* genome assembly (Rhie et al., 2021) (GenBank accession no. [GCA_013347855.1](https://www.ncbi.nlm.nih.gov/datasets/genome/GCA_013347855.1/)). The mapped reads were converted from SAM to BAM files, sorted and indexed using SAMtools v.1.9 (Li et al., 2009b). Variant calling across all 8 individuals was performed using BCFtools v.1.18 (Li, 2011; Li et al., 2009b) functions *mpileup* and *call* with a minimum mapping quality threshold of 20. Initial filtering of the VCF was performed using VCFutils.pl (Li et al., 2009b) and VCFtools v.0.1.16 (Danecek et al., 2011). Only biallelic SNPs called for all individuals, with a minimum variant quality of 20 and combined depth of coverage between 90 and 230, were kept. The coverage thresholds were determined based on inspection of the SNP coverage distribution (Figure S2). A total of 36,458,857 SNPs remained after filtering. Finally, to mitigate background linkage disequilibrium, the filtered VCF was thinned using VCFtools so that no two sites were within 1000 bp of eachother, retaining 870,765 SNPs. Files in bed/bim/fam format were generated from the resulting VCF using PLINK v.1.9 (Purcell et al., 2007). Clustering of individuals was analyzed using ADMIXTURE v.1.3.0 (Alexander et al., 2009) with 10-fold cross-validation and 100 bootstrap replicates for a value of $K=2$.

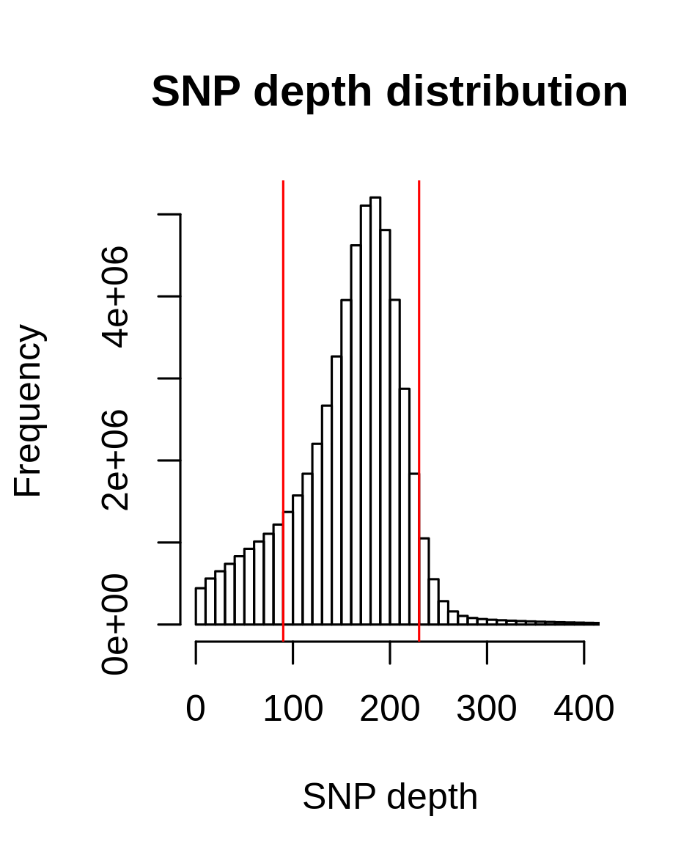

**Figure S2**: Distribution of SNP depth of coverage summed across all eight individuals.

**Results**

The four whole-genome sequences generated in this study showed sequencing depths ranging from 24.71x to 25.87, while the four genome sequences retrieved from SRA showed depths between 18.01x and 23.95x. An overview of mapping statistics for each individual is provided in Table S3. The observed mean nucleotide diversity ($\pi$) was 0.0115 for the two known American eels, 0.0111 for the two known European eels and 0.0120 for the four individuals sampled in Greenland.

| **Individual** | **Number of reads** | **Mapped** | **Properly paired** | **Coverage** | **Insert size** | **HO** |
| --- | --- | --- | --- | --- | --- | --- |
| AM_1 | 208615578 | 0.978 | 0.930 | 18.01 | 499 | 0.01137 |
| AM_2 | 206683547 | 0.977 | 0.932 | 17.72 | 480 | 0.01136 |
| EU_1 | 276832401 | 0.979 | 0.940 | 23.95 | 479 | 0.01100 |
| EU_2 | 241569529 | 0.979 | 0.940 | 20.71 | 477 | 0.01091 |
| L0104 | 179628559 | 0.991 | 0.944 | 25.67 | 254 | 0.01190 |
| L0113 | 179431860 | 0.992 | 0.937 | 25.80 | 245 | 0.01188 |
| L0114 | 179370050 | 0.991 | 0.945 | 25.87 | 258 | 0.01182 |
| L0115 | 179857559 | 0.994 | 0.936 | 24.71 | 230 | 0.01192 |

**Table S3:** Mapping and summary statistics for the eight whole-genome sequenced individuals.

Analyses using ADMIXTURE with $K=2$ clusters clearly separated the known European eels and known Americans eels. The individuals sampled in Greenland clustered with the American eels and exhibited no signs of admixture between the two species, indicating pure American eel ancestry for all four individuals (Fig. S1).

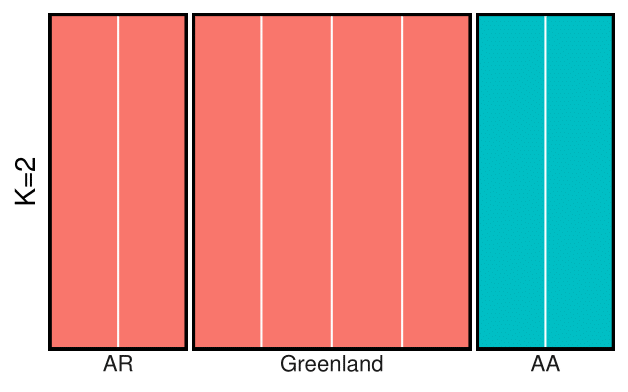

Fig S3. Analyses using ADMIXTURE with $K=2$ clusters clearly separated the known European eels (AA) and known Americans eels (AR). The four individuals sampled in Greenland clustered with the American eels and exhibited no signs of admixture.

Rhie, A., McCarthy, S. A., Fedrigo, O., Damas, J., Formenti, G., Koren, S., Uliano-Silva, M., Chow, W., Fungtammasan, A., Kim, J., Lee, C., Ko, B. J., Chaisson, M., Gedman, G. L., Cantin, L. J., Thibaud-Nissen, F., Haggerty, L., Bista, I., Smith, M., Haase, B., Mountcastle, J., Winkler, S., Paez, S., Howard, J., Vernes, S. C., Lama, T. M., Grutzner, F., Warren, W. C., Balakrishnan, C. N., Burt, D., George, J. M., Biegler, M. T., Iorns, D., Digby, A., Eason, D., Robertson, B., Edwards, T., Wilkinson, M., Turner, G., Meyer, A., Kautt, A. F., Franchini, P., Detrich, H. W., Svardal, H., Wagner, M., Naylor, G. J. P., Pippel, M., Malinsky, M., Mooney, M., Simbirsky, M., Hannigan, B. T., Pesout, T., Houck, M., Misuraca, A., Kingan, S. B., Hall, R., Kronenberg, Z., Sović, I., Dunn, C., Ning, Z., Hastie, A., Lee, J., Selvaraj, S., Green, R. E., Putnam, N. H., Gut, I., Ghurye, J., Garrison, E., Sims, Y., Collins, J., Pelan, S., Torrance, J., Tracey, A., Wood, J., Dagnew, R. E., Guan, D., London, S. E., Clayton, D. F., Mello, C. V., Friedrich, S. R., Lovell, P. V., Osipova, E., Al-Ajli, F. O., Secomandi, S., Kim, H., Theofanopoulou, C., Hiller, M., Zhou, Y., Harris, R. S., Makova, K. D., Medvedev, P., Hoffman, J., Masterson, P., Clark, K., Martin, F., Howe, K., Flicek, P., Walenz, B. P., Kwak, W., Clawson, H., Diekhans, M., Nassar, L., Paten, B., Kraus, R. H. S., Crawford, A. J., Gilbert, M. T. P., Zhang, G., Venkatesh, B., Murphy, R. W., Koepfli, K.-P., Shapiro, B., Johnson, W. E., Di Palma, F., Marques-Bonet, T., Teeling, E. C., Warnow, T., Graves, J. M., Ryder, O. A., Haussler, D., O’Brien, S. J., Korlach, J., Lewin, H. A., Howe, K., Myers, E. W., Durbin, R., Phillippy, A. M., & Jarvis, E. D. (2021). Towards complete and error-free genome assemblies of all vertebrate species. *Nature,* ***592***(7856), 737-746. <https://doi.org/10.1038/s41586-021-03451-0>

**Note S4**

Genotyping was done using 2 loci: mtDNA (cytochrome b)^1^ and 18SrDNA^2^. The polymerase chain reactions (PCR) were performed using Chelex 100^®^ DNA extracts (Bio-Rad Laboratories)^2^ and an amplification profile consisting of a denaturation for 1 min at 94°C, followed by 37 cycles of 30s at 94 C, 30s annealing and elongation at 72 C, followed by 5min at 72°C for final extension. Amplification was carried out using 0.25U of Moltaq DNA Polymerase (Molzym) in 10µl reactions containing 1 µl DNA isolate, 1 µl of 10x PCR buffer, 3mM MgCl2, 250 µM of each deoxyribonucleoside triphosphate and 10 pmol of each primer. Mitochondrial cytochrome b was amplified using the primers CytbF1 5'-CCCTAGTGGATCTACCAACCCC-3', CytbR2seq3 5'-GGGTAGAATAGGGCAAGTATTGTTAG-3', annealing temperature 65°C and 60s elongation. The 18S rRNA gene fragment (412 bp) was amplified using universal primers (SsuF and 18R399, 60 C annealing and 40s elongation)^3,4^.

Categorisation of hybrid individuals was done by means of two additional nuclear gene fragments: blue- and green-sensitive rhodopsin (dso, fwo). The partial gene for the deep sea form of rhodopsin (971 bp, annealing 60°C and 80s elongation) was amplified using the primers opsinfw76 5'-TACCCACAGTACTACCTAGC-3' and opsinrev1046 5'-ACAGAGGACACTGAGGAG-3' (internal opsinrev729 5'-GGTAGTCTCGGACTCCTG-3') which were obtained from an alignment of sequences with the accession numbers L78008, AJ249203, AB043818 and S82619^5^. The fwo fragment (826 bp, annealing 60 C and 80s elongation) was amplified using fwo_fw 5‘-GCAGAACCATGGGCTTATTCAGCT-3‘ and fwo_rev 5‘-TGTAGATGAGGGGGTTGTAGAGG-3‘ based on the reference sequences with accession numbers L78007 and AJ249202^6^.

PCR products (dso) from one putative hybrid were cloned into pGEM-T Vector (Promega) and transformed in Escherichia coli strain TG1^7^. Plasmid DNA was isolated from 16 E. coli colonies per sample using the GFX micro plasmid prep kit (Amersham) and analysed by sequencing. PCR products were purified using Innuprep Gel Extraction Kit (Analytik Jena, Germany) according to the manufacturer’s instructions, and directly sequenced on a CEQ 8000 sequencer (Beckman Coulter GmbH, Krefeld, Germany) using the CEQ Dye Terminator cycle sequencing quick start kit (Beckman Coulter). Sequences were analysed using the software CEQ2000XL (Beckman Coulter), visually edited and aligned using the ClustalW algorithm implemented in BioEdit Sequence Alignment Editor^8^.

1. Tagliavini J, Harrison IJ & Gandolfi G (1995) Discrimination between *A. anguilla* and *A. rostrata* by polymerase chain reaction-restriction fragment length polymorphism analysis. *Journal of Fish Biology*, **47**, 741-743
2. Walsh PS, Metzger DA & Higuchi R (1991) Chelex 100^®^ as a medium for simple extraction of DNA for PCR-based typing from forensic material. *BioTechniques*, **10**, 506-513
3. Englisch U & Koenemann S (2001) Preliminary phylogenetic analysis of selected subterranean amphipod crustaceans, using small subunit rDNA gene sequences. Organisms *Diversity & Evolution*, **1**, 139-145
4. Struck T, Hessling R & Purschke G (2002) The phylogenetic position of the Aeolosomatidae and Parergodrilidae, two enigmatic oligochaete-like taxa of the ‘Polychaeta’, based on molecular data from 18SrDNA sequences. *Journal of Zoological Systematics and Evolutionary Research*, **40**, 155–163
5. Sevilla R. G., Diez A., Norén M., Mouchel O., Jérôme M., Verrez-Bagnis V., et al. (2007). Primers and polymerase chain reaction conditions for DNA barcoding teleost fish based on the mitochondrial cytochrome b and nuclear rhodopsin genes. *Mol. Ecol. Notes*, **7**, 730–734.
6. Zhang,H., Futami,K., Horie,N., Okamura,A., Utoh,T., Mikawa,N.,Yamada,Y., Tanaka,S. and Okamoto,N. (2000) Molecular cloning of fresh water and deep-sea rod opsin genes from Japanese eel Anguilla japonica and expressional analyses during sexual maturation. *FEBS Lett*. **469** (1), 39-43.
7. Sambrook J, Fritsch EF, Maniatis T (1989) Molecular Cloning: A Laboratory Manual, 2nd edn. Cold Spring Harbor Laboratory Press, New York.
8. Hall, T.A. (1999) BioEdit: A User-Friendly Biological Sequence Alignment Editor and Analysis Program for Windows 95/98/NT. *Nucleic Acids Symposium Series*, **41**, 95-98.

**Note S5**

Table S4. Table showing all locations and technical replicates (water samples) which showed qPCR reaction inhibition using 8µL DNA extract.

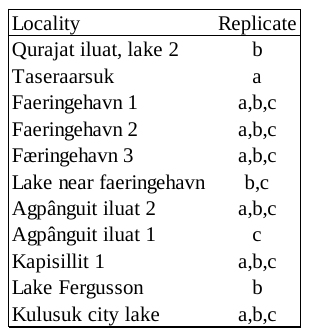

**Note S6**

All sequences have been deposited on Genbank and their corresponding accession numbers can be found in the table below. The below table

**Table S2. Information on all caught eel specimens sampled in Greenland between 2011-2022. NCBI accession numbers and sequence deposit references are included.**

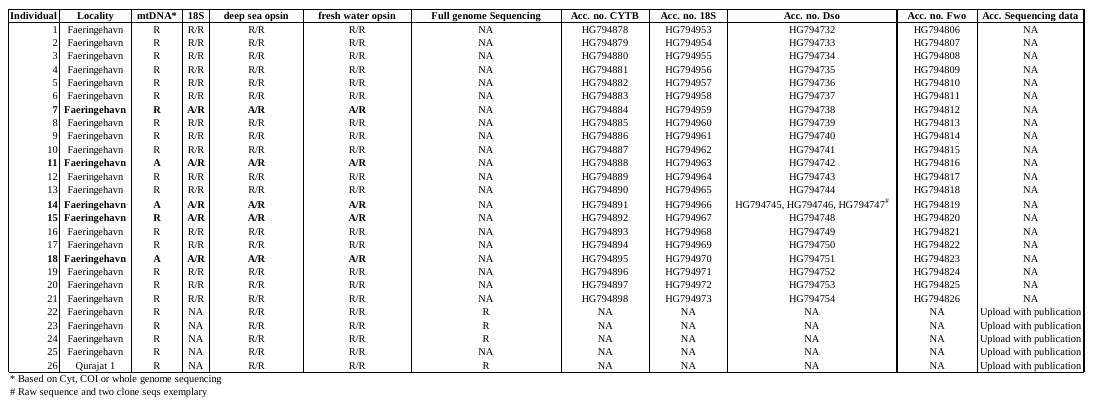
